# Predicting speech-in-noise ability with static and dynamic auditory figure-ground analysis using structural equation modelling

**DOI:** 10.1101/2024.09.08.611859

**Authors:** Xiaoxuan Guo, Ester Benzaquén, Emma Holmes, Joel I Berger, Inga Brühl, William Sedley, Steven P Rushton, Timothy D Griffiths

## Abstract

Auditory figure-ground paradigms assess the ability to extract a foreground figure from a random background, a crucial part of central hearing. Previous studies have shown that the ability to extract static figures (with fixed frequencies) predicts real-life listening: speech-in-noise ability. In this study we assessed both fixed and dynamic figures: the latter comprised component frequencies that vary over time like natural speech. 159 participants (aged 18-79) with a range of peripheral hearing sensitivity were studied. We used hierarchal linear regression and structural equation modelling to examine how well speech-in-noise ability (for words and sentences) could be predicted by age, peripheral hearing, and static and dynamic figure-ground. Regression demonstrated that in addition to the audiogram and age, the low-frequency dynamic figure-ground accounted for significant variance of speech-in-noise, higher than the static figure-ground. The structural models showed that a combination of all types of figure-ground tasks predicted speech-in-noise with a higher effect size than the audiogram or age. Age influenced word perception in noise directly but sentence perception indirectly via effects on peripheral and central hearing. Overall, this study demonstrates that dynamic figure-ground explains more variance of real-life listening than static figure-ground, and the combination of both predicts real-life listening better than hearing sensitivity or age.

## Introduction

Tracking a target sound in a complex auditory scene is one of the core tasks that the auditory system performs and forms an important part of hearing ability. Complaints about understanding speech in noisy environments are frequently encountered in audiology clinics but are difficult to assess because the pure-tone audiogram does not fully reflect this ability (Merten et al., 2022; Besser et al., 2015; George et al., 2007). Sentences-in-noise and word-in-noise tests have been developed to simulate real-life speech-in-noise (SIN) situations, but responses are influenced by other factors, such as levels of education, accent, and language experience as much as hearing and sound segregation. This has motivated work to develop non-speech measures of figure-ground analysis. A non-speech auditory target-in-noise task has been developed called the stochastic figure-ground test (SFG) or fixed-frequency auditory figure ground (AFG-Fixed, Teki et al., 2013). Modelling suggests that sound segregation is achieved based on temporal coherence of the figure (Teki et al., 2013), requiring auditory cortex. Brain studies implicate a network including high-level auditory cortex in humans (Teki et al., 2016; Teki & Griffiths, 2016) and in a primate model (Schneider et al., 2018). Holmes & Griffiths (2019) demonstrated that an AFG discrimination task requiring detection of a gap in auditory figure predicted SIN performance and explained variance in SIN independent of that explained by the pure-tone audiogram (PTA).

SIN perception recruits acoustic features to better segregate sounds in noise. One of the key features is the fundamental frequency corresponding to the perception of pitch in sentences. In this study we assess a type of AFG task in which the frequency components vary over time following the pitch contour of natural speech. This makes the stimulus more speech-like, whilst retaining the overall advantage of the AFG task as a ‘pure’ measure of grouping relevant to real-life listening without linguistic confounds. Two previous studies have investigated AFG with changing frequency patterns (Holmes & Griffiths, 2019; O’Sullivan et al., 2015). Holmes & Griffiths (2019) generated figures that followed the formants of spoken stimuli. While figures generated from the first three formants of speech did not significantly predict SIN, a stimulus based on the first formant and additional components that changed over time coherently with the first formant did predict SIN. We assess similar stimuli here with multiple frequency components that change coherently based on the pitch contours of speech sentences, rather than the frequency of the first formant. We test how well the detection of these frequency components predicts SIN perception.

Natural voiced speech contains multiple harmonics related to the fundamental frequency and is associated with pitch. Harmonicity aids hearing in noise (McPherson et al., 2022). Pitch contributes to SIN processing, especially for people with higher language or hearing competence (Llanos et al., 2021; Shen & Souza, 2018; Huang et al., 2017). In this study, we generated figures related to the harmonic structure of speech, in contrast to the non-harmonic figures used in previous work. We extracted the fundamental frequency from naturally spoken sentences and developed a new type of dynamic auditory figure-ground stimulus using harmonic complexes based on this. We call this the dynamic figure-ground stimulus (AFG-Dynamic). The harmonic features make the auditory figure-ground more speech-like from an acoustic perspective, without incorporating high-level linguistic cues.

We created harmonic complexes in different frequency ranges to explore the importance of the frequency range of the figure. Previous studies suggest that high-frequency hearing sensitivity based on the audiogram may be an important determinant of SIN ability (Polspoel et al., 2022; Zadeh et al., 2019; Holmes & Griffiths, 2019), but have not examined complex figures in different frequency ranges. We constrained the frequency range of the AFG-Dynamic stimuli to low-frequency AFG (AFG -Low) and high-frequency AFG (AFG-High) components to explore how grouping ability in different frequency ranges contributes to SIN perception.

### Predictive measures of Speech-in-Noise Perception

The first aim of the study was to investigate if the new dynamic auditory figure-ground tests are predictive of SIN measures. We hypothesised that both versions of AFG-Dynamic tests (AFG-Low and AFG-High) can predict SiN perception and explain extra variance of SIN independent of the PTA or the prototypical AFG-Fixed. As speech is dynamic in its frequency profile whereas single words have relatively static frequency pattern, we predicted that the fixed-frequency AFG-Fixed better predicts word-level segregation whereas the AFG-Dynamic tests with the changing pitch pattern better predicts sentence-level sound segregation. Three metrics that reflected the SIN ability were used as the outcome measures, including a word-in-noise test (WiN, Guo et al., 2024; Geller et al., 2021), a sentence-in-babble test (SiB, Holmes & Griffiths, 2019), and a subjective self-report measure (The Speech, Spatial and Qualities of Hearing Scale, ‘SSQ’; Gatehouse & Noble, 2004). To capture the word-level perception, the British-ITCP test was used in this study, which has been demonstrated to have a close association with sentence-in-babble performance (Geller et al., 2021). Sentence-level perception is an important aspect of SIN perception and is most commonly assessed as a measure of functional hearing. SiB tests have long been developed to capture patient speech perception in noise (Elliott, 1995; Nilsson et al., 1994; Killion et al., 2004; Wilson et al., 2012). The sentence-in-babble test used here consists of the Oldenburg sentences and babble noise and has been used previously as an assessment of SIN perception (Holmes & Griffiths, 2019). We further included a subjective measure (SSQ) to incorporate participant experience of real-life listening to the overall assessment of SIN ability as well.

### Modelling the Relationships Among Auditory Figure-Ground Perception, Age, and Speech-in-Noise Perception

The second aim of the study was to describe the relationships among the psychoacoustic measures used in the study and identify the contribution of different factors to SIN perception in a multivariate model using structural equation modelling (SEM). The current study had a complex design investigating different measures of hearing thresholds, auditory figure-ground, and speech-in-noise. This type of design favours the use of SEM compared to regression as it allows having multiple observed variables indicating one latent variable (hypothetical constructs that are not directly measured but can be inferred by their observed variables) and reflects the relative importance of indirect effects, such as the interaction between covariates on outcomes. Three conceptual models based on different outcome variables were therefore constructed with assumptions on the direction of causality according to existing literature. The three outcome variables were: the word-in-noise measure, the sentence-in-babble measure, and the two measures combined. As the word- and sentence-level SIN analysis and the self-reported SIN ability tap into different domains of SIN perception, models predicting the three SIN measures separately should provide additional information on the differences of the three domains of SIN analysis when interacting with AFG, PTA and Age. We also investigated the domain-general SIN by combining the three measures into one latent variable.

The fixed-frequency AFG test has been shown to predict SIN perception (Holmes & Griffiths, 2019). In this study, we added the two additional dynamic AFG measures with high and low frequencies to form an AFG latent variable that predicts SIN perception.

In terms of the exogenous variables, PTA and the participant’s age were taken into account. PTA has been shown to predict SIN ability (George et al., 2007; Wong et al., 2008; Besser et al., 2015; Bochner et al., 2015; Holmes & Griffiths, 2019). Age has also been recognised as a key factor impacting both hearing and SIN perception (see Billings & Madsen, 2018 for a review on the topic). Deterioration of the auditory periphery – including hair cell and cochlear nerve loss, as well as cochlear synaptopathy (Dias et al., 2024; Liu et al., 2024; Xie et al., 2024) could all lead to decreased real-life listening ability, and these peripheral deteriorations are all tied with aging (Chadha et al., 2021). Holmes and Griffiths^8^ found a relationship between age and both AFG and SIN performance. Altered auditory peripheral function could result in lowered frequency and temporal resolution, which would inevitably impact the central sound segregation ability measured by the AFG tests. However, the relationships between AFG and SIN perception retained after accounting for age and PTA, which indicates that figure-ground measures may also index PTA- and age-independent deficits in SIN perception. Thus, we hypothesised that PTA and Age would predict both SIN and AFG with Age impacting PTA, and that AFG can independently predict SIN after accounting for Age and PTA.

## Methods

### Participants

A total of 170 participants were recruited, of whom 11 were excluded due to data quality as per criteria described later. The final sample size used for analysis was 159. The sample had a wide range of age (mean = 45.95, SD = 18.47, range = 18 – 79) as well as hearing thresholds (mean = 13.30 dB, SD = 9.90 dB, see Figure 1 for more detailed audiogram results), with 105 female participants. All participants were neurotypical native English speakers with no history of auditory disorders, no history of speech and language disorders, and who were not currently taking any psychotropic drugs. Informed consent was obtained from participants before the experiments. The study was approved by the Newcastle University Ethics Committee (46225/2023).

**Figure 1.**
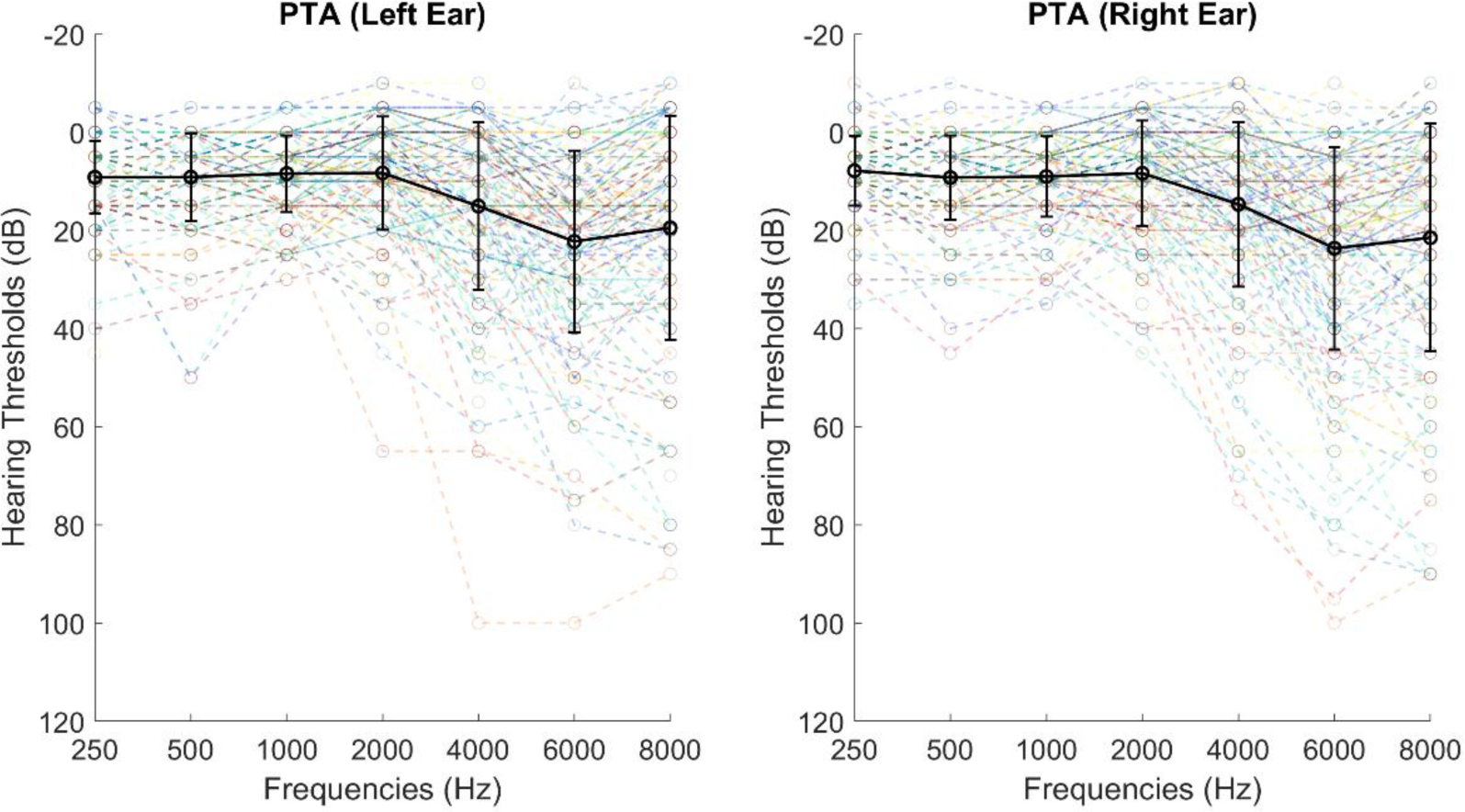
The distribution of hearing sensitivity of 250 – 8000 Hz for the left and the right ear separately of all participants. The x axis shows the frequencies measured, and the y axis shows hearing thresholds measured in decibels. The coloured lines with circles plot individual audiogram results, and the thicker black line with circles and error bars show the averaged group PTA. The error bars display the standard deviation.

### Stimuli and Tasks

#### Fixed-Frequency Auditory Figure-Ground Gap Discrimination Task

The AFG-Fixed gap discrimination test was the same as that developed by Holmes & Griffiths (2019); an auditory figure was composed of temporally coherent pure-tone elements (each 50 ms duration) repeating over time. Each figure was 42 chords long with 3 figure components per chord (i.e. coherence level of 3). The figure was superimposed on an auditory ground, which is a tone cloud made of pure-tone elements (also 50 ms duration each) of randomised (or stochastic) frequencies between 180 – 7246 Hz. In each trial, two figure-ground stimuli were presented to the participants, sequentially with an inter-stimulus interval of 400 ms. A gap (6 chords long) was present in either figure. Although, importantly, the ground tones continued through the gap, so participants needed to have segregated the figure from the ground to perform this task. The participants were instructed to choose which of the two figure-ground stimuli contained a gap in the figure. The test used a 1-up 1-down adaptive procedure, starting at signal-to-noise ratio (SNR) of 6 dB and varied systematically across trials. The step size started at 2 dB and went down to 0.5 dB after 3 reversals. Two runs were presented to each participant with different exemplars, with both runs terminating after 10 reversals. The median of the last 6 reversals for both runs were taken and averaged as a measure of performance. Higher SNR scores indicate worse performance.

#### Dynamic Auditory Figure-Ground Pattern Discrimination Task

In contrast to the prototype AFG-Fixed that has a fixed-frequency pattern over time, the novel dynamic AFG contains pitch information akin to speech. The pitch contours were extracted from the English Oldenburg sentences read by a male British speaker (Holmes & Griffiths, 2019), using Praat version 6.2.09 with a time step of 0.75/75 Hz (100 pitch values per second), and had a frequency range of 74.94 – 295.44 Hz (M=131.59, SD=15.61). The low-frequency noise (below 10 Hz) and artificial high frequencies (above 300 Hz) introduced by the Praat periodicity analysis were removed to obtain pitch trajectories (see Figure 2(a) for an example). There are gaps in natural pitch tracks as shown in Figure 2 (a). To avoid the participants using these gaps, the natural speech gaps and stops were first removed from the pitch contour. An example of the conjoined signal is shown in Figure 2(b). As demonstrated in the plot, the new signal has a general downwards trend and some sharp transitions caused by the removal of the gaps and linear interpolation. To remove the drift from individual signals, we demeaned the signal and applied detrending to the demeaned signal. Low-pass filtering (minimum-order filter with a stopband attenuation of 60 dB) with a 2000 Hz cutoff frequency was then carried out to remove the artificial spikes. The trend and the mean were then added back to the filtered signal to keep the final signal as similar to the original pitch trajectory as possible. An example of the final pitch signal is plotted in red in Figure 2(b).

**Figure 2.**
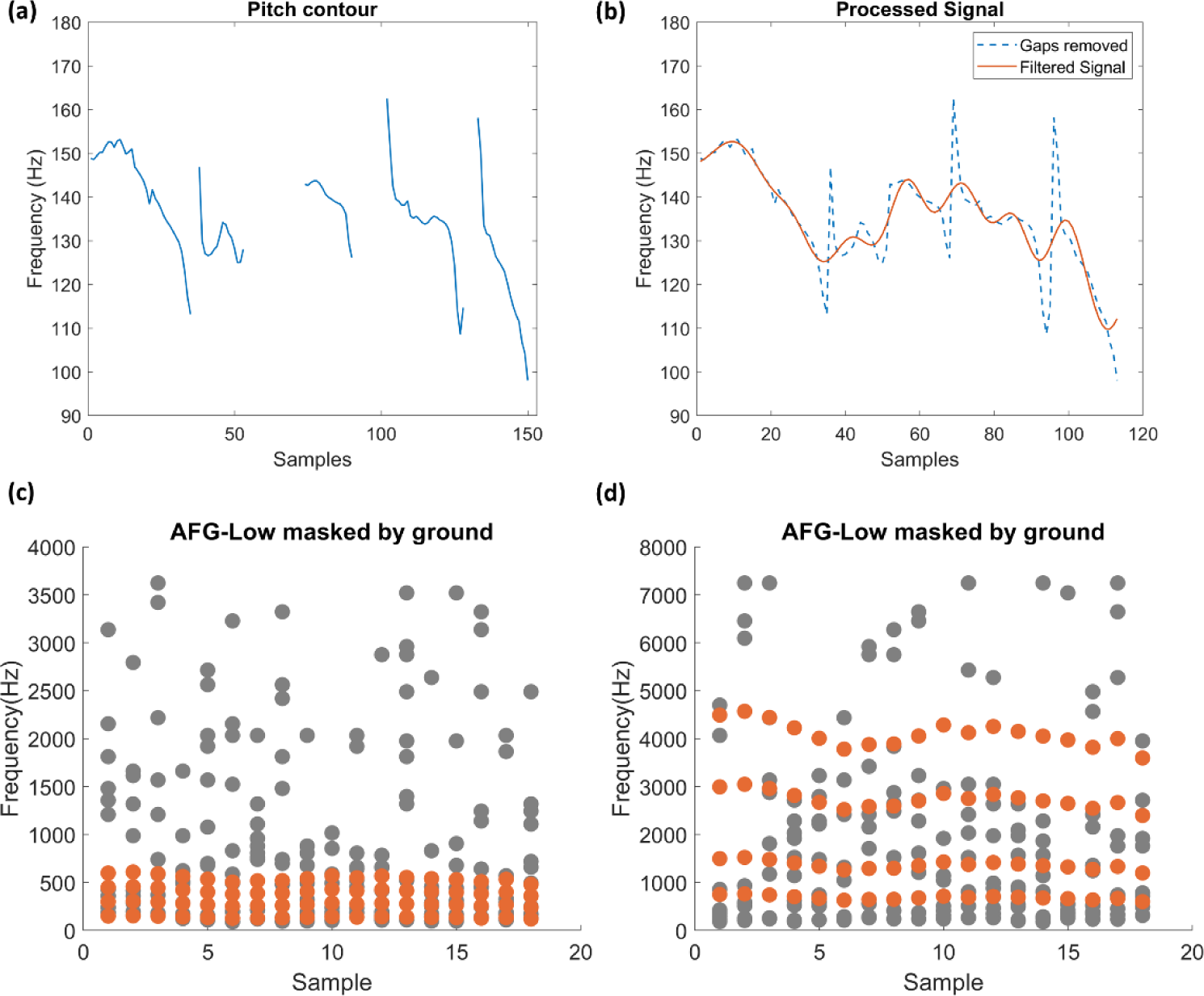
The figure shows the extraction of pitch contour (Figure 2(a) (b)) and the AFG stimuli with the pitch contour embedded (Figure 2(c) (d)). Figure 2(a) and (b) shows that the pitch information extracted from the sentence “Alan brought four small desks”. The x axis plots the time in seconds and the y axis shows the frequencies in Hz. Figure 2 (a) shows the raw pitch contour plotted against time. Figure 2 (b) shows the pitch trajectory after being processed. The blue line is the pitch contour with the gaps removed. The red line shows the final processed signal. The dotted plots illustrate examples of the two different types of AFG-Dynamic stimuli. Figure 2 (c) shows the lower-frequency dynamic AFG. Figure 2 (d) on the right side is the high-frequency dynamic AFG. The x axis shows the time in milliseconds and the y axis shows the frequency in Hz. Figure elements are depicted in orange while ground elements are depicted in grey.

After processing the pitch signals, the resultant frequency profiles were grouped into 50-ms long segments by computing the average to form the figure elements. The second, third, and fourth harmonics of these fundamental components were used to construct the remaining elements of each chord (see Figure 2(c)) for the AFG-Low. The tones were gated with a 10 ms raised-cosine ramp to smooth the onset and offset of the sounds. The high-frequency figure (AFG-High) retained the pitch trajectories of the low-frequency version, but the components were the fundamental frequencies multiplied by 5, 10, 20, and 30. The top frequency of each figure was checked not to exceed the masking frequencies. Like the AFG-Fixed stimuli, the auditory ground was comprised of randomised pure-tone segments on a logarithmic scale. Although, while ground tones for the AFG-High stimuli used the same range of frequencies as the AFG-Fixed (180 Hz–7246 Hz), AFG-Low stimuli used ground tones with a lower frequency range (90 – 3623 Hz; in other words, half of the upper and lower frequency values from the AFG-High stimuli) to achieve a better masking effect. See Figure 2 (c) (d) for an illustrated example of the two types of stimuli.

The two AFG-Dynamic tests (AFG-Low and AFG-High) had the same paradigm and were counter-balanced across participants. Within each test, both the figure and the ground stimuli were presented every trial, either with the same or different figure pattern. In case of a trial with different figure patterns, the durations of the figures were matched but the frequency elements were based on different pitch trajectories. The ground elements were tailored to different figures. The tests used a two-alternative forced-choice task, which required the participants to hold the sounds in memory and decide whether or not the second figure had the same pattern as the first one. The inter-stimulus (within each trial) interval was 0.2 seconds. A two-down one-up staircase procedure was used with a total of 22 reversals. The initial SNR was 12 dB with a step size of 2 dB, which then changed to 0.5 dB after 7 reversals. The final score was calculated by averaging the dB SNR of the last 6 reversals and a higher SNR would indicate poorer performance. The sound pairs, which were matched for duration, and the trial orders were kept the same across participants. The same design was used for both the low-frequency and the high-frequency versions of the AFG-Dynamic test.

### Speech-in-Noise

#### Word-in-Noise Test

The WiN test was the British Iowa Test of Consonant Perception (B-ITCP) adapted from the Iowa Test of Consonant Perception ((Guo et al., 2024)). Target speech sounds were monosyllabic words with a consonant-vowel-consonant (CVC) or CVCC structure. The test used British accents including one female and one male voice. The babble noise was an 8-talker babble, presented at -2 dB signal-to-noise ratio (SNR). The onset of the auditory target was between 0.5 and 1.0 s, randomly positioned from the babble onset. The babble segment of each trial was randomly selected from a 15-seconds babble stimulus. The length of the words varied from 0.304 – 0.757 s (mean: 0.508 s, SD: 0.086 s). Participants were asked to choose the word they heard out of a list of 4 words displayed on the screen balanced for difficulty and phonetic contrasts. Percent correct across trials was taken as the outcome measure for the WiN test. This is the only test that was scored differently, and a higher score for the WiN test indicates better performance.

#### Sentence-in-Babble Test

Details of the SiB test are reported in Holmes & Griffiths (2019). The target sentences were English Oldenburg sentences consisting of 5 words following a structure of Name-Verb-Number-Adjective-Noun (e.g., “Alan brought four small desks”). The sentences were masked by 16-talker babble. The target sentences appeared 500 ms after babble onset and ended 500 ms before babble offset. Participants were shown a 5*10 matrix on the screen, where each word in the sentence had 10 options. The test used a one-down one-up adaptive paradigm with the starting SNR at 0 dB. The total number of reversals was 10 and the step size began at 2 dB and decreased to 0.5 dB after 3 reversals. The task had two runs interleaved. The target sentences were different in each run. The final score was calculated by averaging the dB SNRs of the last 6 reversals across the 2 runs. A lower score in this test indicates better performance.

#### Speech, Spatial and Qualities of Hearing Scale

The subjective self-report SIN ability was assessed using the Speech, Spatial and Qualities of Hearing Scale-speech hearing (SSQ, Gatehouse & Noble, 2004). Two of the questions were removed from the shortened speech-hearing questionnaire due to their ambiguity. See Appendix 1 for the full list of questions used in this questionnaire. Each item has a score from 0 to 10 with the higher score indicating more difficulty in hearing.

#### Procedure

An audiometry test was carried out, followed by the 5 computer tasks, which were presented in a fixed order for all participants, except that the order of the AFG-High and the AFG -Low tests were counterbalanced across participants. The tasks were presented in the following order: (1) SiB, (2) AFG-Dynamic test (AFG -High or AFG-Low, determined by counterbalancing across participants), (3) SSQ, (4) WiN, (5) second version of the AFG-Dynamic test (AFG-High or AFG -Low, whichever they had not already completed), (6) AFG-Fixed. Participants were asked to sit in the front of a computer monitor (Dell Inc.) used to present the tasks. The auditory stimuli were presented through headphones (Sennheiser HD 380 Pro) linked to a sound card (RME FireFace UC).

### Data Analysis

#### Test of Correlation for AFG-Dynamic

The outcome measures of SiB and the AFG tests were the medians of the last 6 reversals. The performance was considered stable if the performance differences of the last 6 reversals were smaller than ± 5 dB. Participants who did not show stable performance were excluded from the final analysis. Bivariate correlations and hierarchical regressions were carried out to explore the relationship between AFG-Dynamic and SIN tests. Tests of normality (Kolmogorov-Smirnov) showed that AFG-High and WiN were not normally distributed, so Spearman’s rho was used to examine the hypotheses regarding the relationships between the three speech measures with AFG-Dynamic (low and high version), AFG-Fixed, PTA and age. Holm-Bonferroni correction was applied to correct for multiple comparisons. As linear regression is a more tolerant measure for non-normality, stepwise regression was conducted to check if there were predictive relationships between SIN and AFG as well as specifying the variance explained by individual predictors. The order of entering the independent variables is based on how close their associations were according to the correlation results (Figure 4). These tests were performed using SPSS version 28 and visualised with MATLAB R2021a.

**Figure 3.**
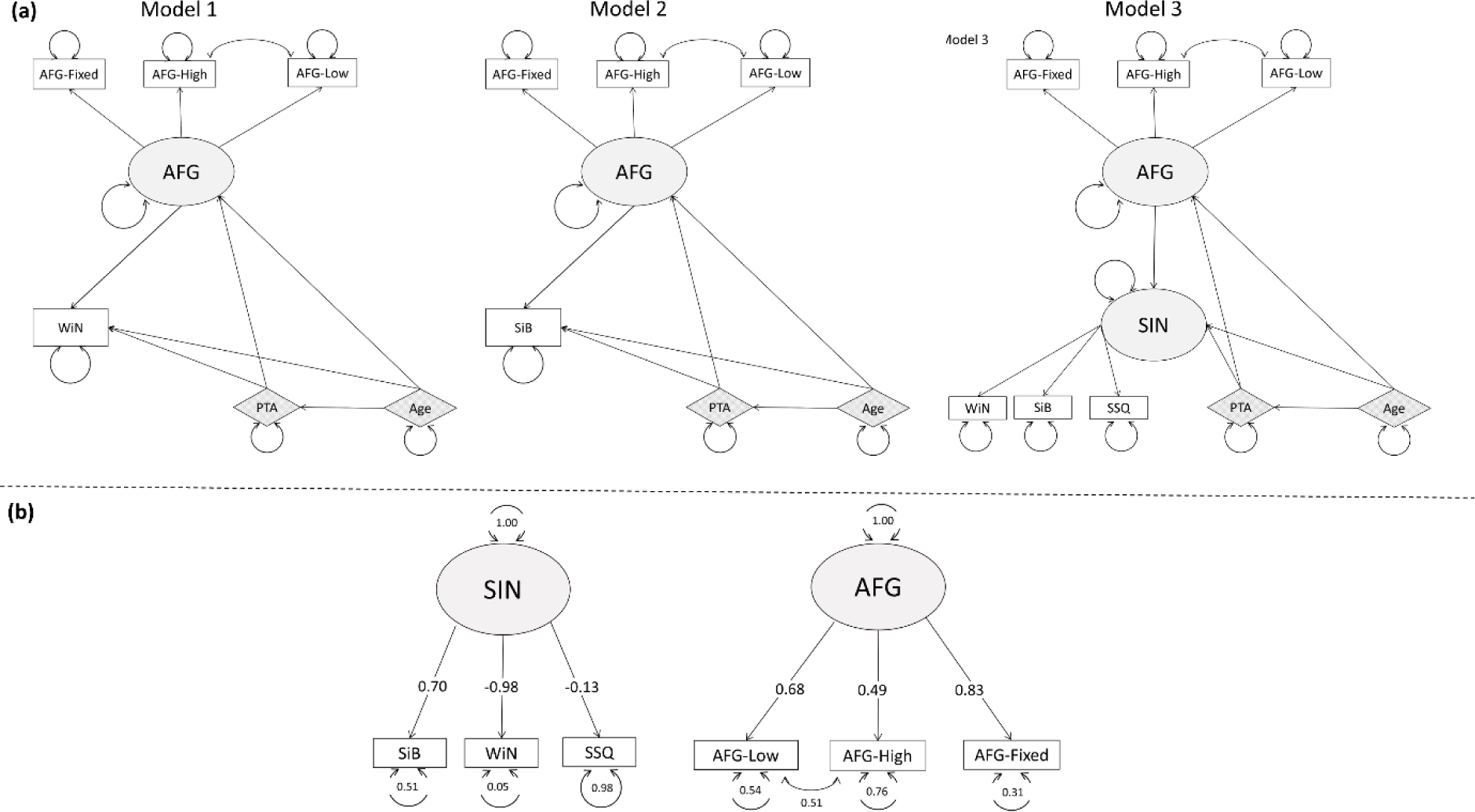
Figure 3 (a) shows the conceptual models of WiN(Model 1), SiB perception (Model 2), and SIN with combined word and sentence perception (Model 3). The shaded ovals represent latent variables, the rectangles represent observed variables, and the diamonds with striped shading are exogenous variables. The arrowed circle of each variable represents the error (the size of the circle is not proportional to the radius). The indicators have arrows pointing to them from the latent variables. External variables point the arrows to the latent variables to suggest a causal effect on the latent variables. Figure 3 (b) shows the CFA of SIN and AFG. Shaded ovals represent the latent variables, rectangular boxes are the indicators, and the circles associated with each variable are the residuals. Latent variables are connected to their indicators through arrows pointing to the indicators. The error for SIN and AFG is 1 as they are not subject to any causal influences in this limited model.

**Figure 4.**
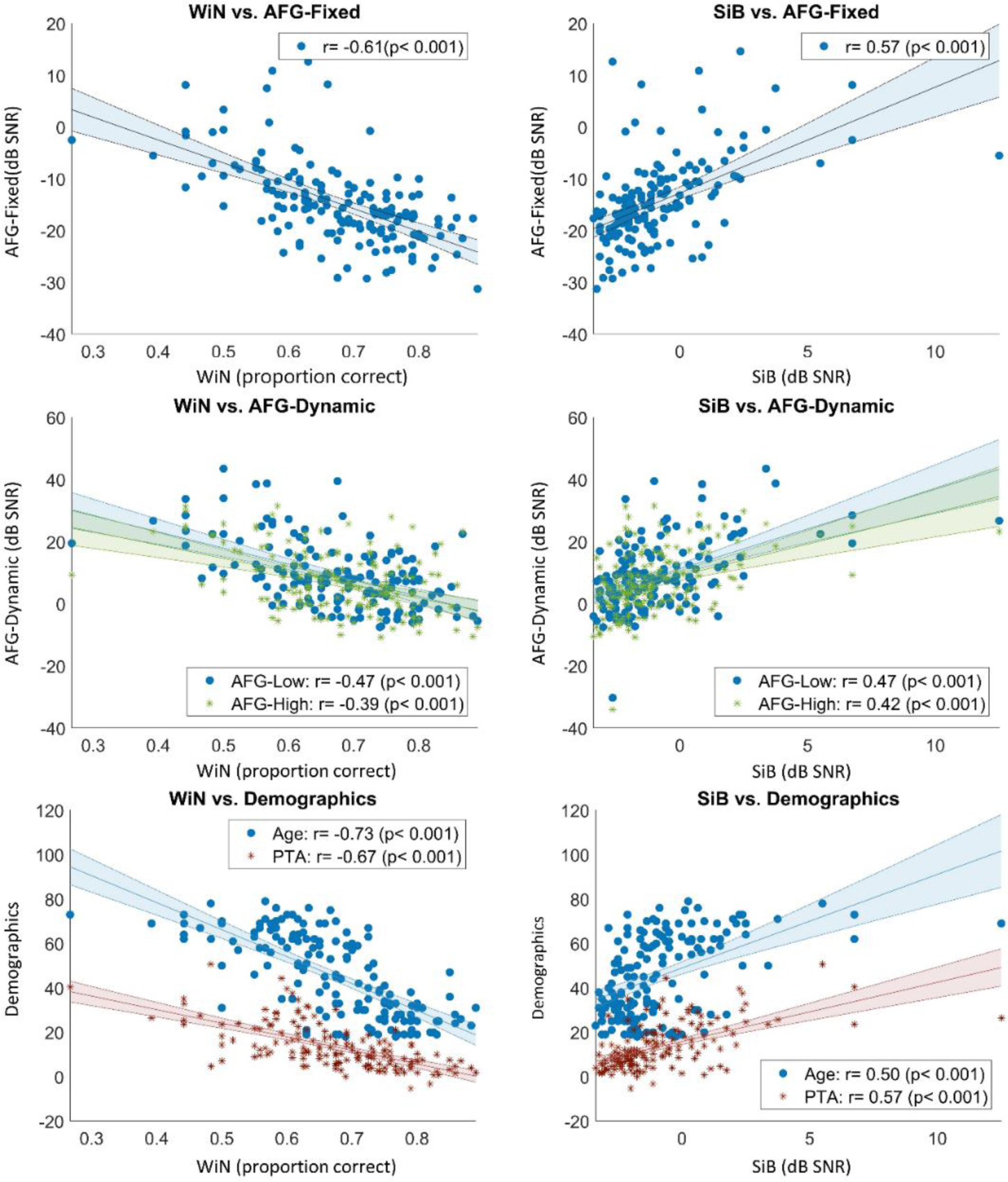
Bivariate correlation among variables. The p values shown here were corrected. The lines of best fit were plotted as straight lines in the figure with shaded error bars. The x axis for the left panel showed the WiN results as proportion correct (number of correct answers divided by the overall number of trials), and the y axes of the figures on the left panel from top to down were the two AFG tasks measured in dB SNR and Age and PTA. On the right panel were three figures of the same variables plotted against the SiB test results measured in dB SNR.

#### Modelling the Inter-Relations of Predictors of SIN

To account for the inter-relationships of the indicator variables, structural equation modelling (SEM) was conducted using the lavaan (version 0.6-15) package in R (version 4.2.1). Maximum likelihood estimation was used with nonnormality correction based on the Satorra-Bentler scaled test statistic. Robust measures were reported in this study (Brosseau-Liard et al., 2012; Brosseau-Liard & Savalei, 2014).

Initial conceptual models (Model 1 and 2) are illustrated in Figure 3 (a). Model 1&2 were devised to explore word-level and sentence-level SIN analysis separately. Model 3 illustrates a combined model of all three SIN measures. In all three models, the SIN measures were predicted by AFG indicated by the AFG-Fixed, and two AFG-Dynamic measures. PTA and Age also predicted SIN as well as AFG. To decide the latent variable structure, a confirmatory factor analysis (CFA) was used to examine the measurement quality on a subset of the data (101 participants) before conducting the final analysis. Figure 3 (b) demonstrates the CFA models of the two latent constructs in the three models: SIN and AFG. While there are no rules of thumb defining the acceptable thresholds of a factor loading, SSQ as a measure of functional hearing should predict a large variance of SIN tests similar to the other two SIN indicators. The SSQ however had a visibly weak connection to SIN and thus was removed from further analysis. All three AFG indicators seemed to be acceptable to be entered to the final model.

The results of this analysis were used to guide the selection of scaling variable (Bollen et al., 2022). Scaling variables are used to assign scales to latent variables, which is essential when identifying a model. The method used in lavaan is the Fixed Marker (FM) scaling that fixes the loading of the chosen scaling variable to 1(*Lavaan.Org - Model Syntax 2*, n.d.). The path coefficients (abbreviated as β) can be interpreted as: one SD of variable A increase leads to a β SD increase of variable B while all other relevant connections were held constant. The residual or measurement error of the indicators represents variance unexplained by the measure “due to random measurement error, or score unreliability” (Kline, 2015, pp9).

The WiN measure was chosen as the scaling variable based on its high path coefficient connecting to the SIN latent variable. WiN was the only test measured by percentage, which resulted in a difference in scale of the outcome compared to the other tests. This was re-scaled via z-scoring (removing the mean and dividing the results by the standard deviation (SD) of the original scores of WiN). Importantly, contrary to the measures assessed with SNR, a higher score of WiN indicated better performance. This means one SD increase from the mean in WiN would lead to a β SD decrease in SIN. However, since WiN was used as the scaling variable, the SIN latent variable took the scale of WiN, SiB instead showed a negative path coefficient. The different interpretation of SNR- and percent correct -based scoring would further influence other factors connecting to SIN. To avoid confusion and simplify results interpretation, the WiN results were multiplied by -1 so a higher score would indicate worse performance.

AFG-High, AFG-Low, and AFG-Fixed are the indicators of the latent variable AFG. Similarly, the AFG-Fixed was chosen as the scaling variable due to its close connection with the AFG latent variable. AFG-High and AFG-Low were made with similar parameters except for the frequency range and should tap into very similar mechanisms, hence they covary. The three SEM models further consisted of Age and PTA as exogenous variables, which were both configured to predict SIN and AFG.

The model fit was evaluated by a set of criteria (Hu & Bentler, 1999; Kline, 2015). These includes the Bentler comparative fit index (CFI) and Tucker-Lewis Index (TLI), the root-mean-square error of approximation (RMSEA), the standardised root mean squared residual (SRMR), and the chi-square test. Both RMSEA and SRMR are absolute measures of the estimated discrepancy between the predicted and observed models. The SRMR is a measure of the mean absolute correlation residual measuring the differences between the original correlations (observed) and the implied correlations by the model. RMSEA ≤ 0.06 and SRMR ≤ 0.08 have been suggested to indicate a close model fit (Hu & Bentler, 1999). RMSEA up to 0.10 is considered fair fit, but above 0.10 is generally unacceptable (Browne & Cudeck, 1992). CFI and TLI, on the other hand, are incremental indices that reflect the relative improvement of the model fit compared to a baseline model (Kline, 2015).

TLI is non-normed so it can fall outside the 0-1 range whereas CFI is normed, but the cutoff for both of them is above 0.95 for a good fit (Hu & Bentler, 1999). The chi-square (χ^2^) result was also reported (Kline, 2015). The null hypothesis for the chi-square test is that the predicted model reflects the true data perfectly. Thus, a nonsignificant chi-square would indicate a good model fit.

### Results

The descriptive statistics are reported in Table 1. WiN had the narrowest range of data distribution (Table 1), with an accuracy of around 67% correct. All other tests were measured in signal-to-noise ratio, with the AFG tests showing a wider spread of performance than SiB (Table 1). The two AFG-Dynamic tests had positive thresholds whereas the fixed-frequency AFG elicited negative thresholds, indicating that the AFG-Dynamic tests were more challenging.

**Table 1.** The mean and standard deviation of the participants’ performance on the five computer tasks.

|  | Mean | Standard Deviation |
| --- | --- | --- |
| SiB | -0.880 | 2.114 |
| WiN | 0.673 | 0.107 |
| AFG-Low | 8.991 | 10.897 |
| AFG-High | 7.252 | 10.447 |
| AFG-Fixed | -14.542 | 8.200 |

#### Relationships between SIN measures and AFG-Dynamic

Both the sentence-level and the word-level SIN tests showed moderate to strong correlations with all the AFG measures, hearing, and age (Figure 4). After correction, all p-values remained highly significant. SSQ, however, did not show any significant correlation with other speech measures (p > 0.34) and was removed from further analysis.

The hierarchical regression predicting SiB performance gave three significant predictors, revealing that PTA, AFG-Low, and AFG-Fixed performance significantly predicted SiB performance (F (3, 155) = 39.879, p < .001). The model accounted for 43.56 % of the variance in SiB. Table 2 specifies the variance explained by the significant predictors. For the WiN model, four significant predictors were significant: age, PTA, AFG-Low, and AFG-Fixed (F (4, 154) = 62.560, p < .001). The model accounted for 61.90% of the variance in WiN. Table 2 specifies the variance explained by each predictor. For SiB, PTA was the best predictor explaining about 31% of the model with the AFG-Low adding 9.9% to the model. Whereas, for WiN, Age seemed to be the strongest predictor. Of the significant predictors, AFG-Fixed added the least variance to both SiB and WiN, which was about 1%∼2% after accounting for the other variables.

**Table 2.** This table displays the R² values and p values of models including an increasing number of predictors that add significant variance to the models predicting either SiB or WiN.

| SiB | Standardis<br>ed<br>Coefficients<br>Beta | R<sup>2</sup> | p | WiN | Standardis<br>ed<br>Coefficients<br>Beta | R<sup>2</sup> | p |
| --- | --- | --- | --- | --- | --- | --- | --- |
| PTA | 0.368 | 0.314 | < 0.001 | Age | -0.409 | 0.498 | < 0.001 |
| + AFG-Low | 0.253 | 0.413 | < 0.001 | +AFG-Low | -0.217 | 0.580 | < 0.001 |
| + AFG-Fixed | 0.197 | 0.436 | 0.015 | + PTA | -0.218 | 0.609 | 0.003 |
| +AFG-High | 0.126 | - | 0.125 | + AFG-Fixed | -0.134 | 0.619 | 0.049 |
| +Age | 0.074 | - | 0.400 | +AFG-High | -0.121 | - | 0.075 |

#### Structural Equation Model of SIN, AFG, Hearing, and Age

The fit indices for the three models are shown in Table 3, and path coefficients are plotted in Figure 5. All fit indices for Model 1 and Model 2 were within our criteria. The path coefficients in Model 1 were all significant. Model 2 had mostly significant paths with a nonsignificant one of Age to SiB.

**Figure 5.**
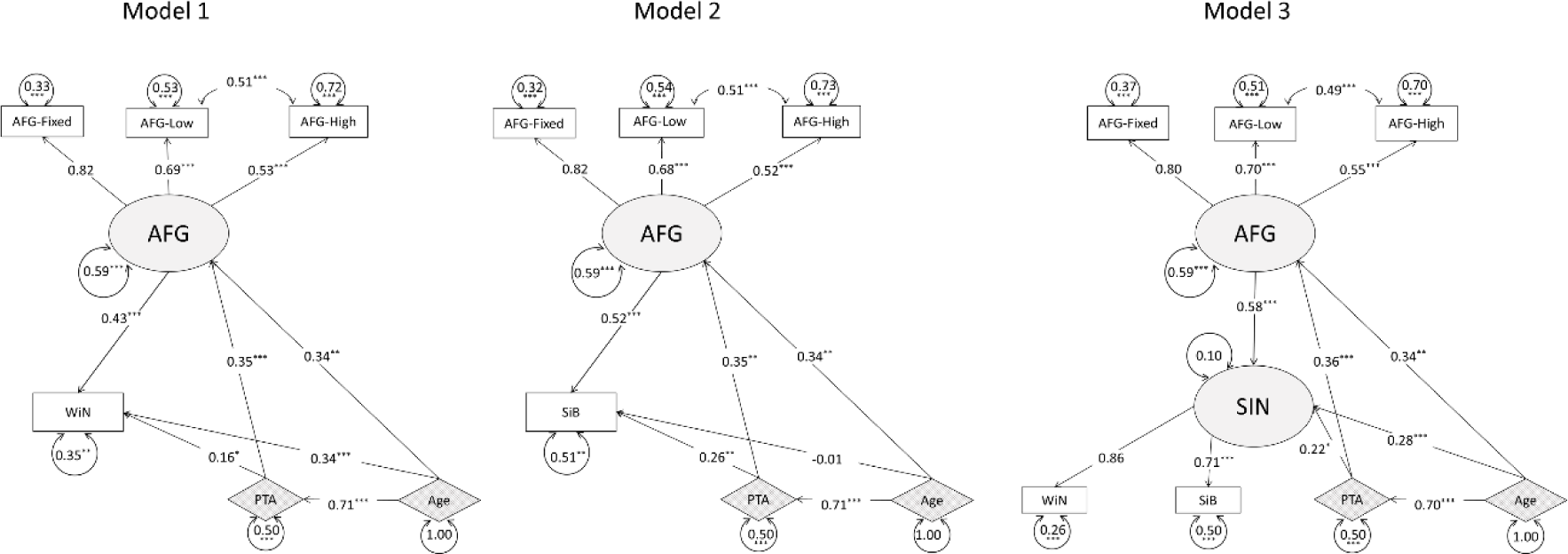
Model 1 and Model 2 are presented with either the WiN measure or the SiB measures as the dependent variable, Model 3 had WiN and SiB combined as the dependent variable. All indicator variables were plotted in rectangular. The oval shape represents the latent variable (AFG) in both models, and the external variables are plotted in diamond shape. The latent variable measured by indicators have arrows pointing towards the indicators. Otherwise, the arrows point from the variable that causes change in another one. The path coefficients were marked by numbers and error terms were marked by both numbers and a circle around the number. The significance level was marked by asterisks. Three asterisks represent p < 0.001, two represent p< 0.01, one represents p< 0.05. Note that while AFG-Fixed in all three models and WiN in Model 3 were not marked with asterisks, it was not because they failed to predict the latent variables but because the scaling variables were not estimated in the SEM. The scaling variables dominate the latent constructs.

**Table 3.** Fit indices for Model 1, 2 and 3.

| Fit Index | Model 1 | Model 2 | Model 3 |
| --- | --- | --- | --- |
| $\chi^2$ | 9.547 (p=0.92) | 7.910 (p=0.91) | 20.617 (p=0.940) |
| RMSEA | 0.079 | 0.065 | 0.092 |
| CFI | 0.990 | 0.992 | 0.978 |
| TLI | 0.969 | 0.975 | 0.949 |
| SRMR | 0.028 | 0.029 | 0.036 |

As expected, in both Models 1 & 2, all three AFG indicators showed significant contributions to the AFG indicator, and the AFG-High and AFG-Low shared significant covariance. AFG-Fixed contributed to AFG with the largest path coefficient (|β| = 0.82) followed by the two dynamic AFG measures. The latent AFG variable predicted WiN and SiB significantly, with the largest variance compared to PTA and Age in both models. PTA had a significant but smaller contribution to each SIN measure. Age only has a significant direct impact on WiN and not on SiB.

Model 3 followed the conceptual model structure shown in Figure 3 but had the path connecting SSQ to SIN removed as it was not significant. This model met most of the set criteria for an excellent model fit except for the RMSEA (Table 3). RMSEA incorporates model complexity and models with smaller degrees of freedom tend to obtain a poorer RMSEA (Kenny et al., 2015). This pattern of results is similar to that obtained for Models 1 and 2, which also had excellent fit based on most of the indicators but poorer than expected RMSEA. However, as the combined results of other indicators all showed that the model fits the data very well, we deem that this model is acceptable.

Similar to Models 1 and 2, AFG explained the largest variance of the latent SIN variable (β = 0.56) in Model 3 (Figure 5), compared to Age and PTA. Both SiB and WiN showed significant contributions to the latent SIN variable, but WiN had a numerically greater contribution (|β| = 0.86 for WiN compared to |β| = 0.71 for SiB). Age was the second largest predictor of SIN, after AFG, and PTA had a smaller (but nevertheless significant) direct impact on SIN. Although, both PTA and Age had a significant indirect impact on SIN through AFG.

## Discussion

### Predicting SIN Perception with Dynamic AFG in the Linear Models

This study showed a moderate to strong correlation between all AFG measures and SIN, both on the word and the sentence level. The low-frequency AFG came out as a significant predictor of WiN and SiB, explaining the largest variance in both models after accounting for demographic factors (Age or PTA). It is unexpected that even for the WiN model the AFG-Low explained more variance than the static AFG. The dynamic AFG was designed to carry the fundamental frequency patterns and so should better predict sentence-level sound segregation than word-level. AFG-Fixed, on the other hand, had no frequency change over time, which was considered more similar to WiN perception. Based on the regression results, however, it seems that adding the speech pitch pattern to the AFG stimuli only improved its predictive power of SIN in general, not specifically to sentence-level perception. This general improvement could be the reason the AFG-Fixed did not explain a higher portion of the variance of SIN as well. Considering that AFG-Low combined both the mechanism of segregating static figure from ground employing the figure’s temporal coherence feature, and speech-like frequency pattern to aid SIN perception, it is reasonable to see higher variance obtained by AFG-Low in a linear model.

One possible explanation for the relationship between AFG-Low and both word and sentence-level SIN is its harmonic structure. AFG-Fixed differed from AFG-Dynamic in two ways: it is both static and inharmonic. Some of the AFG-Fixed stimuli might contain near-harmonic figures by chance, but most of the stimuli were inharmonic, which elicit weaker pitch perception (Micheyl et al., 2012). Pitch plays an important role in SIN perception (Meha-Bettison et al., 2018; Shen & Souza, 2018; Binns & Culling, 2007; Carroll & Zeng, 2007), the mechanism of which was reviewed by Oxenham (2008). This includes not only its strong association with the stress contour of a whole sentence but also other linguistic features such as phonemes in words. The pitch information embedded in the AFG-Low can help with differentiating the envelope fluctuations of the target sound from the background sound, which is key for speech intelligibility. Thus, the stronger pitch strength could be the reason that AFG-Low, while sharing the same basic principles with the static AFG, predicts SiB or WiN better.

The high-frequency dynamic AFG had a numerically weaker correlation as was hypothesised and did not explain additional variance in WiN or SiB after accounting for other tasks. This could be because AFG-High shared a high covariance with AFG-Low due to the similar parameters used for these two tests. The AFG-Low more closely resembles the speech stimuli used in this study with its frequency range being closely configured to the pitch range of speech formants, which might be the reason that AFG-Low outperformed AFG-High in predicting SIN.

The SSQ measure did not correlate with either of the speech measures, which was not a unique finding (Oberfeld & Klöckner-Nowotny, 2016; Ertürk et al., 2023). This could be because the shorter SSQ version does not have enough sensitivity to capture SIN perception as only a few questions were related to speech comprehension in human speech noise.

### Predicting SIN Perception in a Multivariate Model

The linear regression models displayed the core contribution of the new dynamic AFG measure as well as the static measure. However, the stepwise procedure did not account for the interaction between variables. The SEM model provided a more comprehensive picture of the experiment that went beyond ranking the important predictors of SIN measures.

Firstly, Models 1 and 2 showed that all three AFG predictors have an impact on the SIN performance. This means that when accounting for the interaction and covariance shared between the indicators, all of the AFG predictors should be considered a necessary part of the auditory figure-ground analysis. Interestingly, while the linear measures showed a tighter relationship between AFG-Low and WiN/SiB, AFG-Fixed in the SEM models contributes the most to the AFG latent variable. This suggest that as the ‘prototype’ AFG, the static AFG that assesses people’s ability to pick up the temporally coherent figure from the tone cloud, is still the core of the AFG analysis process. Combined with the regression results, it shows that the dynamic pitch pattern does add an important aspect to AFG and should be used in combination with AFG-Fixed. Based on their individual predictive value of the regression results, in a linear model, when using both measures are not possible, AFG-Low should be a better test to measure SIN ability compared to AFG-Fixed.

The combined AFG measures explained the largest variance (43%, 52%) of both speech measures in Model 1 & 2, compared to Age and PTA. This also differs from the linear regression results, where PTA or Age was found to be the greatest predictor. This difference suggests that AFG tasks have greater predictive power in combination as various ways to assess scene segregation, whereas each AFG task individually assesses slightly different abilities that are weaker individually than the influence of the demographic factors. The lower path coefficient of PTA compared to AFG indicates that the ability to process speech (either single-word utterances or sentences) in a noisy environment directly relies more on segregating auditory streams and tracking the pattern of the target sounds over time than simply picking up acoustic signals as measured by PTA. However, PTA also had a mediation effect on WiN/SiB through a large path coefficient to AFG. This means that in addition to a relatively small direct impact on SIN perception, deteriorated peripheral hearing could alter functional hearing through modifying central sound processing, which is consistent with our hypothesis.

A mediation effect was also evident with the Age-driven impact on SIN perception. Age led to 71% SD change in PTA in this study, meaning the PTA variance was largely dominated by age-related hearing loss. Age also decreased central sound processing measured by AFG here by 34%, consistent with the results reported by Holmes and Griffiths (2019). However, while Age showed a significant correlation with SiB, it did not modify SiB performance directly in the SEM model, which is consistent with the regression results. WiN is a harder task for people who are older or with higher hearing thresholds. This is because less in the way of compensatory mechanisms can be employed for hearing a short word compared to a sentence that contains a legitimate syntactic structure. While normal ageing can result in deteriorated hearing sensitivity and the perception of other acoustic properties (fine structure or harmonicity), language perception skills are generally preserved (Burke & Mackay, 1997).

The interaction among predictors in Model 3 did not change much after combining the WiN and SiB into one latent variable. WiN and SiB had a similar level of contribution to the SIN latent factor and the small residual term of SIN suggests that WiN and SiB together provide a holistic measurement of SIN, with a small effect of unmeasured sources of unique variance on the latent variable. It is important to highlight however, that combining the measures into a latent variable could hide the different effects of other predictors such as Age, like in Models 1 and 2.

### Limitations and Future Direction

The sample size of the current study might have caused the fit to be suboptimal. There is no golden rule in terms of determining an appropriate sample size for SEM. Researchers have suggested a variety of standards based on the number of observations (N) per statistical estimates (q) ranging from 20:1 to 5:1 depending on the complexity of the model (Bentler & Chou, 1987; Kline, 2015) or an absolute sample size of 250 if using the Satorra-Bentler scaled method (Hu & Bentler, 1999). The current study has around 8:1 N:q, which is sufficient to find a good solution to meet the convergence criteria, but not optimal. Further studies are needed to validate the model with a larger sample size.

This study also focussed mainly on individuals without symptomatic hearing impairment. The new dynamic measures will need to be tested on different populations such as hearing impaired or patients with cochlear implants, to see if the results can be replicated with people who struggle with SIN perception. Indeed, recent data suggest that AFG-fixed does predict SIN performance in CI users (Choi et al., 2023), so it is plausible that AFG-dynamic in CI users may explain even further variance. This then can potentially be used for clinical practice to assess patients’ dynamic sound segregation. Future research can also incorporate other aspects of SIN perception (e.g., subcortical sound analysis, language ability) and cognitive measures (general intelligence, auditory working memory) to test if the effect of AFG on SIN holds when accounting for these other factors.

In conclusion, these data show that an adequate model of SIN perception needs to account for age, peripheral auditory function, and measures of grouping that we have previously demonstrated to have a brain basis. We introduced new measures of central grouping in this work that incorporate harmonicity and a pitch trajectory taken from natural speech. These measures have improved the prediction of speech in noise in the multivariate model.

## Supporting information

Appendix The SSQ questionnaire

## Author contributions

Xiaoxuan Guo: conceptualisation, study design, data collection, data analysis, manuscript writing

Ester Benzaquén: conceptualisation, study design

Emma Holmes: study design

Joel I Berger: study design

Inga Brühl: data collection

William Sedley: conceptualisation, study design

Steven Rushton (co-senior author): data analysis

Timothy D Griffiths (co-senior author): conceptualisation, study design, data analysis, funding.

All authors have edited the manuscript and approved its submission to this journal.

## Data availability

Data will be available upon request.

## Additional Information

This work was supported by the MRC [grant number MR/T032553/1]. There are no conflicts of interest, financial, or otherwise.

## References

Bentler, P. M., & Chou, C.-P. (1987). Practical Issues in Structural Modeling. https://journals.sagepub.com/doi/10.1177/0049124187016001004

Besser, J., Festen, J. M., Goverts, S. T., Kramer, S. E., & Pichora-Fuller, M. K. (2015). Speech-in-speech listening on the LiSN-S test by older adults with good audiograms depends on cognition and hearing acuity at high frequencies. Ear and Hearing, 36(1), 24–41. 10.1097/AUD.0000000000000096

Billings, C., & Madsen, B. (2018). A perspective on brain-behavior relationships and effects of age and hearing using speech-in-noise stimuli. Hearing Research, 369. 10.1016/j.heares.2018.03.024

Binns, C., & Culling, J. F. (2007). The role of fundamental frequency contours in the perception of speech against interfering speech. The Journal of the Acoustical Society of America, 122(3), 1765–1776. 10.1121/1.2751394

Bochner, J. H., Garrison, W. M., & Doherty, K. A. (2015). The NTID speech recognition test: NSRT(®). International Journal of Audiology, 54(7), 490–498. 10.3109/14992027.2014.991976

Bollen, K. A., Lilly, A. G., & Luo, L. (2022). Selecting scaling indicators in structural equation models (sems). Psychological Methods. 10.1037/met0000530

Brosseau-Liard, P. E., & Savalei, V. (2014). Adjusting Incremental Fit Indices for Nonnormality. Multivariate Behavioral Research, 49(5), 460–470. 10.1080/00273171.2014.933697

Brosseau-Liard, P. E., Savalei, V., & Li, L. (2012). An Investigation of the Sample Performance of Two Nonnormality Corrections for RMSEA. Multivariate Behavioral Research, 47(6), 904–930. 10.1080/00273171.2012.715252

Browne, M. W., & Cudeck, R. (1992). Alternative Ways of Assessing Model Fit. Sociological Methods & Research, 21(2), 230–258. 10.1177/0049124192021002005

Burke, D. M., & Mackay, D. G. (1997). Memory, language, and ageing. Philosophical Transactions of the Royal Society of London. Series B, Biological Sciences, 352(1363), 1845–1856. 10.1098/rstb.1997.0170

Carroll, J., & Zeng, F.-G. (2007). Fundamental frequency discrimination and speech perception in noise in cochlear implant simulations. Hearing Research, 231(1– 2), 42–53. 10.1016/j.heares.2007.05.004

Chadha, S., Kamenov, K., & Cieza, A. (2021). The world report on hearing, 2021. Bulletin of the World Health Organization, 99(4). 10.2471/BLT.21.285643

Choi, I., Gander, P. E., Berger, J. I., Woo, J., Choy, M. H., Hong, J., Colby, S., McMurray, B., & Griffiths, T. D. (2023). Spectral Grouping of Electrically Encoded Sound Predicts Speech-in-Noise Performance in Cochlear Implantees. Journal of the Association for Research in Otolaryngology: JARO, 24(6), 607–617. 10.1007/s10162-023-00918-x

Dias, J., McClaskey, C., Alvey, A., Lawson, A., Matthews, L., Dubno, J. R., & Harris, K. (2024). Effects of age and noise exposure history on auditory nerve response amplitudes: A systematic review, study, and meta-analysis. Hearing Research, 447. 10.1016/j.heares.2024.109010

Elliott, L. L. (1995). Verbal auditory closure and the speech perception in noise (SPIN) Test. Journal of Speech and Hearing Research, 38(6), 1363–1376. 10.1044/jshr.3806.1363

Ertürk, P., Aslan, F., & Türkyılmaz, M. D. (2023). Listening to speech-in-noise with hearing aids: Do the self-reported outcomes reflect the behavioral speech perception task performance? European Archives of Oto-Rhino-Laryngology. 10.1007/s00405-023-08193-5

Gatehouse, S., & Noble, W. (2004). The Speech, Spatial and Qualities of Hearing Scale (SSQ). International Journal of Audiology, 43(2), 85–99.

Geller, J., Holmes, A., Schwalje, A., Berger, J. I., Gander, P. E., Choi, I., & McMurray, B. (2021). Validation of the Iowa Test of Consonant Perception. The Journal of the Acoustical Society of America, 150(3), 2131–2153. 10.1121/10.0006246

George, E. L. J., Zekveld, A. A., Kramer, S. E., Goverts, S. T., Festen, J. M., & Houtgast, T. (2007). Auditory and nonauditory factors affecting speech reception in noise by older listeners. The Journal of the Acoustical Society of America, 121(4), 2362–2375. 10.1121/1.2642072

Guo, X., Benzaquén, E., Holmes, E., Choi, I., McMurray, B., Bamiou, D.-E., Berger, J. I., & Griffiths, T. D. (2024). British Version of the Iowa Test of Consonant Perception (p. 2024.09.04.611204). bioRxiv. 10.1101/2024.09.04.611204

Holmes, E., & Griffiths, T. D. (2019). ‘Normal’ hearing thresholds and fundamental auditory grouping processes predict difficulties with speech-in-noise perception. Scientific Reports, 9(1), Article 1. 10.1038/s41598-019-53353-5

Hu, L., & Bentler, P. M. (1999). Cutoff criteria for fit indexes in covariance structure analysis: Conventional criteria versus new alternatives. Structural Equation Modeling: A Multidisciplinary Journal, 6(1), 1–55. 10.1080/10705519909540118

Huang, Y. T., Newman, R. S., Catalano, A., & Goupell, M. J. (2017). Using prosody to infer discourse prominence in cochlear-implant users and normal-hearing listeners. Cognition, 166, 184–200. 10.1016/j.cognition.2017.05.029

Kenny, D. A., Kaniskan, B., & McCoach, D. B. (2015). The Performance of RMSEA in Models With Small Degrees of Freedom. Sociological Methods & Research, 44(3), 486–507. 10.1177/0049124114543236

Killion, M. C., Niquette, P. A., Gudmundsen, G. I., Revit, L. J., & Banerjee, S. (2004). Development of a quick speech-in-noise test for measuring signal-to-noise ratio loss in normal-hearing and hearing-impaired listeners. The Journal of the Acoustical Society of America, 116(4 Pt 1), 2395–2405. 10.1121/1.1784440

Kline, R. B. (2015). Principles and Practice of Structural Equation Modeling, Fourth Edition: Fourth Edition (4th edition). Guilford Press.

lavaan.org—Model syntax 2. (n.d.). Retrieved 8 March 2024, from https://lavaan.ugent.be/tutorial/syntax2.html

Liu, J., Stohl, J., & Overath, T. (2024). Hidden hearing loss: Fifteen years at a glance. Hearing Research, 443, 108967. 10.1016/j.heares.2024.108967

Llanos, F., German, J. S., Gnanateja, G. N., & Chandrasekaran, B. (2021). The neural processing of pitch accents in continuous speech. Neuropsychologia, 158, 107883. 10.1016/j.neuropsychologia.2021.107883

McPherson, M. J., Grace, R. C., & McDermott, J. H. (2022). Harmonicity aids hearing in noise. *Attention*, Perception & Psychophysics, 84(3), 1016–1042. 10.3758/s13414-021-02376-0

Meha-Bettison, K., Sharma, M., Ibrahim, R. K., & Mandikal Vasuki, P. R. (2018). Enhanced speech perception in noise and cortical auditory evoked potentials in professional musicians. International Journal of Audiology, 57(1), 40–52. 10.1080/14992027.2017.1380850

Merten, N., Boenniger, M. M., Herholz, S. C., & Breteler, M. M. B. (2022). The Associations of Hearing Sensitivity and Different Cognitive Functions with Perception of Speech-in-Noise. Ear and Hearing, 43(3), 984–992. 10.1097/AUD.0000000000001154

Micheyl, C., Ryan, C. M., & Oxenham, A. J. (2012). Further evidence that fundamental-frequency difference limens measure pitch discrimination. The Journal of the Acoustical Society of America, 131(5), 3989–4001. 10.1121/1.3699253

Nilsson, M., Soli, S. D., & Sullivan, J. A. (1994). Development of the Hearing in Noise Test for the measurement of speech reception thresholds in quiet and in noise. The Journal of the Acoustical Society of America, 95(2), 1085–1099. 10.1121/1.408469

Oberfeld, D., & Klöckner-Nowotny, F. (2016). Individual differences in selective attention predict speech identification at a cocktail party. eLife, 5, e16747. 10.7554/eLife.16747

O’Sullivan, J. A., Shamma, S. A., & Lalor, E. C. (2015). Evidence for Neural Computations of Temporal Coherence in an Auditory Scene and Their Enhancement during Active Listening. The Journal of Neuroscience: The Official Journal of the Society for Neuroscience, 35(18), 7256–7263. 10.1523/JNEUROSCI.4973-14.2015

Oxenham, A. J. (2008). Pitch Perception and Auditory Stream Segregation: Implications for Hearing Loss and Cochlear Implants. Trends in Amplification, 12(4), 316–331. 10.1177/1084713808325881

Schneider, F., Dheerendra, P., Balezeau, F., Ortiz-Rios, M., Kikuchi, Y., Petkov, C. I., Thiele, A., & Griffiths, T. D. (2018). Auditory figure-ground analysis in rostral belt and parabelt of the macaque monkey. Scientific Reports, 8(1), 17948. 10.1038/s41598-018-36903-1

Shen, J., & Souza, P. E. (2018). On Dynamic Pitch Benefit for Speech Recognition in Speech Masker. Frontiers in Psychology, 9, 1967. 10.3389/fpsyg.2018.01967

Teki, S., Barascud, N., Picard, S., Payne, C., Griffiths, T. D., & Chait, M. (2016). Neural Correlates of Auditory Figure-Ground Segregation Based on Temporal Coherence. Cerebral Cortex (New York, N.Y.: 1991), 26(9), 3669–3680. 10.1093/cercor/bhw173

Teki, S., Chait, M., Kumar, S., Shamma, S., & Griffiths, T. D. (2013). Segregation of complex acoustic scenes based on temporal coherence. eLife, 2, e00699. 10.7554/eLife.00699

Teki, S., & Griffiths, T. D. (2016). Brain Bases of Working Memory for Time Intervals in Rhythmic Sequences. Frontiers in Neuroscience, 10, 239. 10.3389/fnins.2016.00239

Wilson, R. H., McArdle, R., Watts, K. L., & Smith, S. L. (2012). The Revised Speech Perception in Noise Test (R-SPIN) in a multiple signal-to-noise ratio paradigm. Journal of the American Academy of Audiology, 23(8), 590–605. 10.3766/jaaa.23.7.9

Wong, L. L. N., Cheung, C., & Wong, E. C. M. (2008). Comparison of hearing thresholds obtained using pure-tone behavioral audiometry, the Cantonese Hearing in Noise Test (CHINT) and cortical evoked response audiometry. Acta Oto-Laryngologica, 128(6), 654–660. 10.1080/00016480701642189

Xie, R., Wang, M., & Zhang, C. (2024). Mechanisms of age-related hearing loss at the auditory nerve central synapses and postsynaptic neurons in the cochlear nucleus. Hearing Research, 442, 108935. 10.1016/j.heares.2023.108935

