## Appendix The SSQ questionnaire for "Predicting speech-in-noise ability with static and dynamic auditory figure-ground analysis using structural equation modelling"

### SSQ: Speech

Participant ID:.....

Date:.....

|  |  |
| --- | --- |
| 1. You are talking with one other person and there is a TV on in the same room. Without turning the TV down, can you follow what the person you're talking to says? | Not at all Perfectly<br>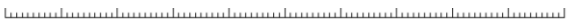<br>0 1 2 3 4 5 6 7 8 9 10<br>Min Max |
| 2. You are talking with one other person in a quiet, carpeted lounge-room. Can you follow what the other person says? | Not at all Perfectly<br>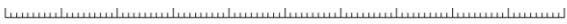<br>0 1 2 3 4 5 6 7 8 9 10<br>Min Max |
| 3. You are in a group of about five people, sitting round a table. It is an otherwise quiet place. You can see everyone else in the group. Can you follow the conversation? | Not at all Perfectly<br>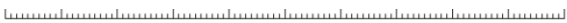<br>0 1 2 3 4 5 6 7 8 9 10<br>Min Max |
| 4. You are in a group of about five people in a busy restaurant. You can see everyone else in the group. Can you follow the conversation? | Not at all Perfectly<br>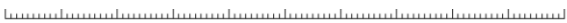<br>0 1 2 3 4 5 6 7 8 9 10<br>Min Max |
| 5. You are talking with one other person. There is continuous background noise, such as a fan or running water. Can you follow what the person says? | Not at all Perfectly<br>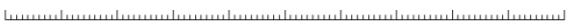<br>0 1 2 3 4 5 6 7 8 9 10<br>Min Max |
| 6. You are in a group of about five people in a busy restaurant. You *cannot* see everyone else in the group. Can you follow the conversation? | Not at all Perfectly<br>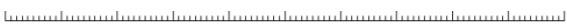<br>0 1 2 3 4 5 6 7 8 9 10<br>Min Max |
| 7. You are talking to someone in a place where there are a lot of echoes, such as a church or railway terminus building. Can you follow what the other person says? | Not at all Perfectly<br>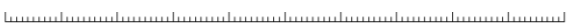<br>0 1 2 3 4 5 6 7 8 9 10<br>Min Max |

|  |  |
| --- | --- |
| 8. You are listening to someone talking to you, while at the same time trying to follow the news on TV. Can you follow what both people are saying? | Not at all<br>Perfectly<br>012345678910<br>MinMax |
| 9. You are in conversation with one person in a room where there are many other people talking. Can you follow what the person you are talking to is saying? | Not at all<br>Perfectly<br>012345678910<br>MinMax |
| 10. You are with a group and the conversation switches from one person to another. Can you easily follow the conversation without missing the start of what each new speaker is saying? | Not at all<br>Perfectly<br>012345678910<br>MinMax |
| 11. Can you easily have a conversation on the telephone? | Not at all<br>Perfectly<br>012345678910<br>MinMax |
| 12. You are listening to someone on the telephone and someone next to you starts talking. Can you follow what's being said by both speakers? | Not at all<br>Perfectly<br>012345678910<br>MinMax |
